## Supplementary figures and images for "Purging viral latency by a bifunctional HSV-vectored therapeutic vaccine in chronically SIV-infected macaques"

### Figure S1.tif

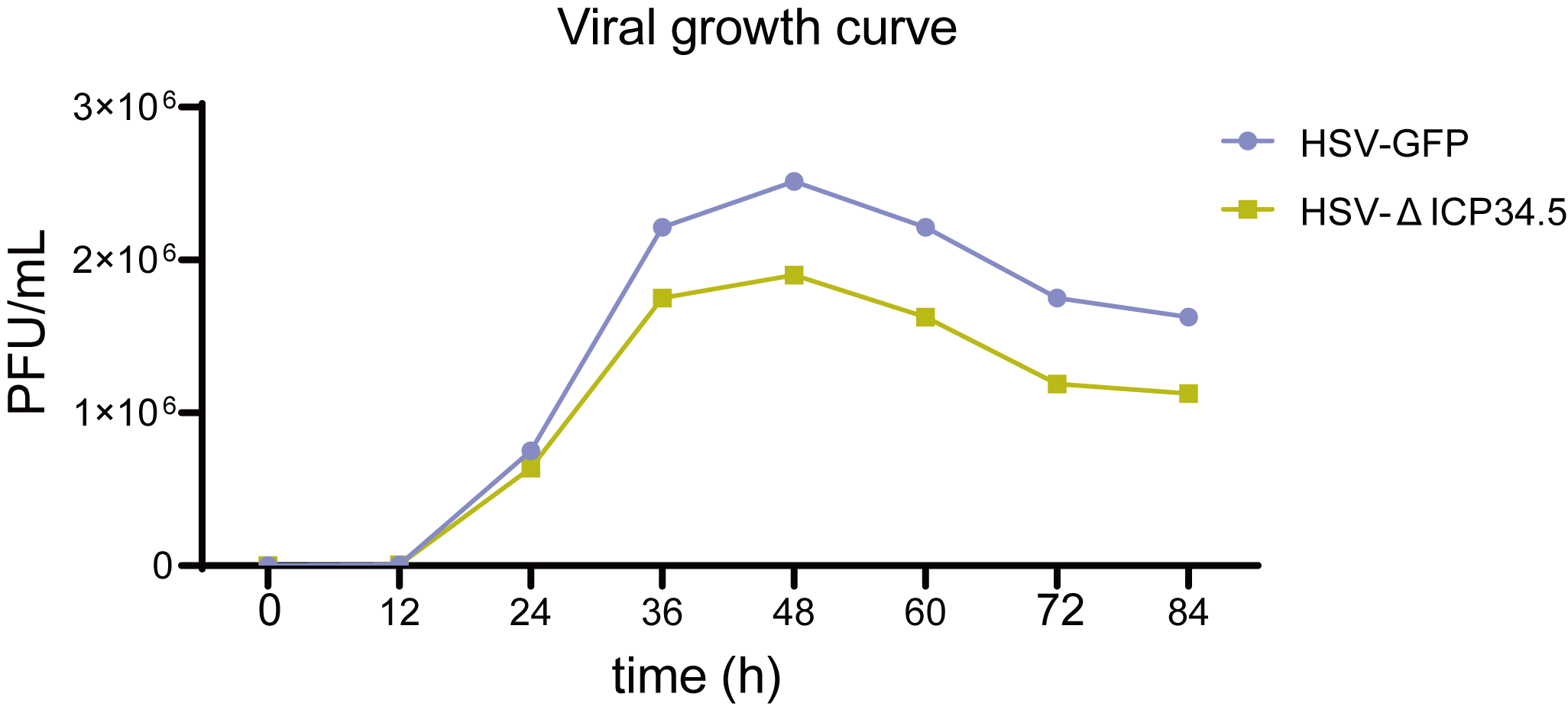

### Figure S2.tif

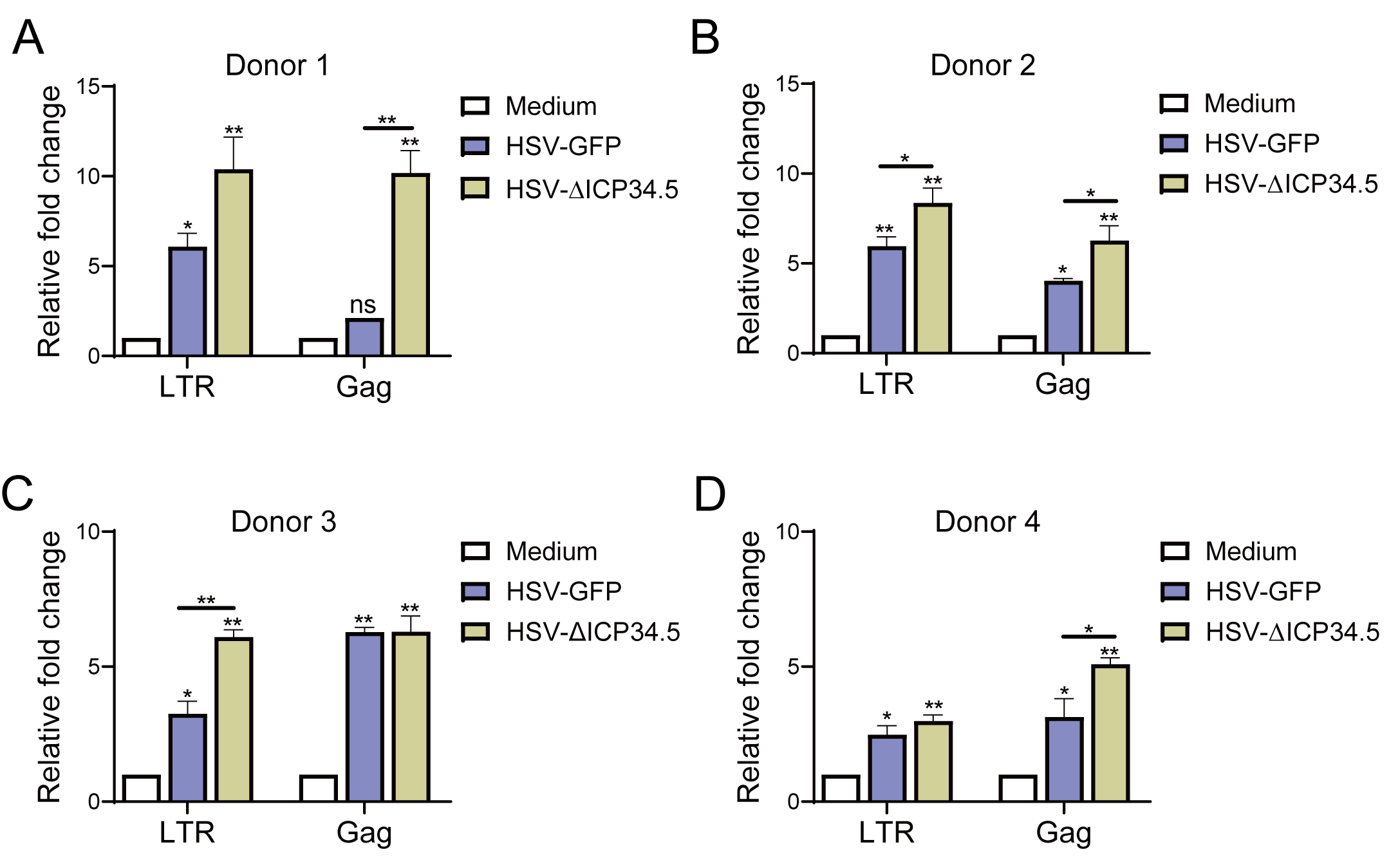

### Figure S3.tif

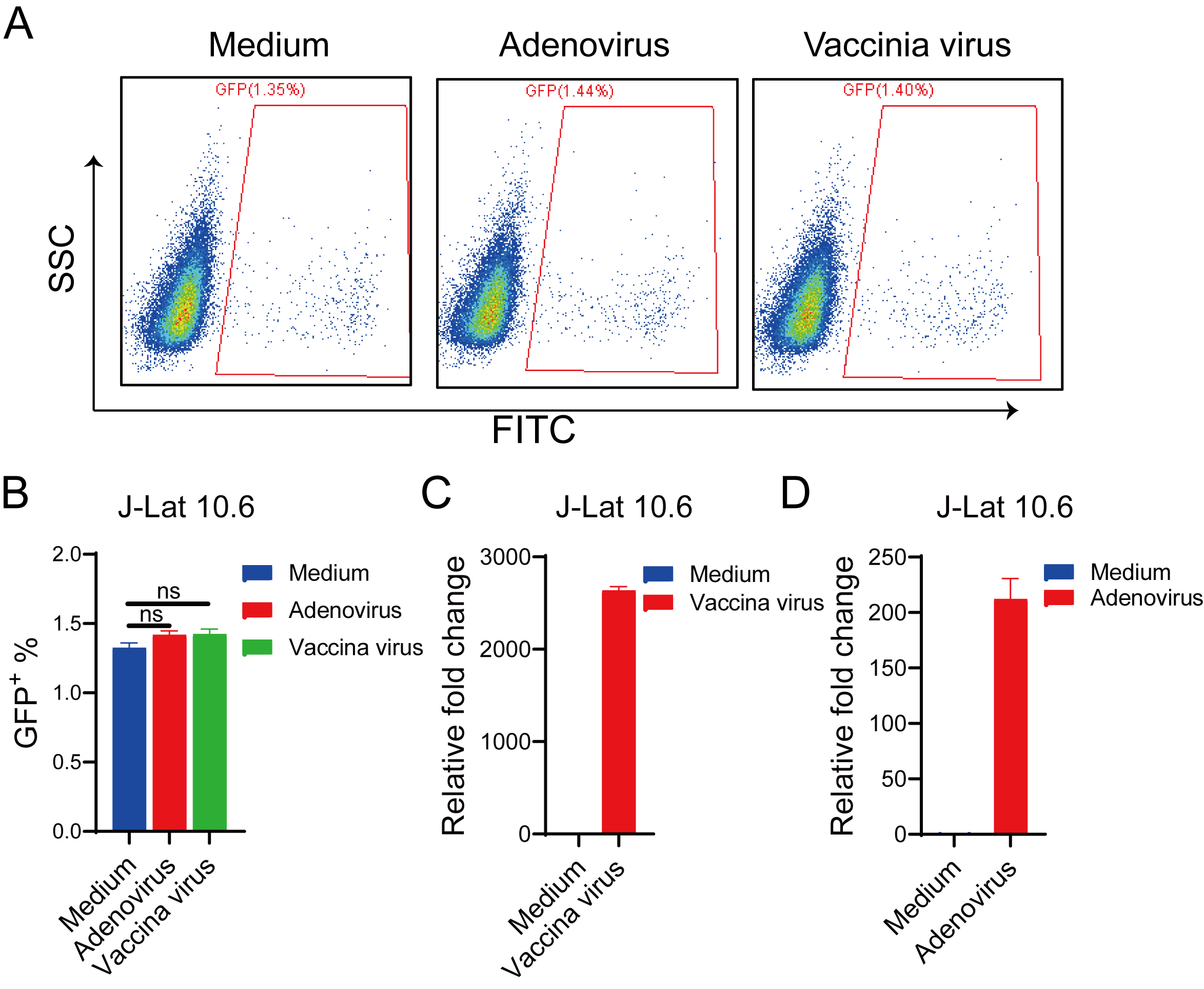

### Figure S4.tif

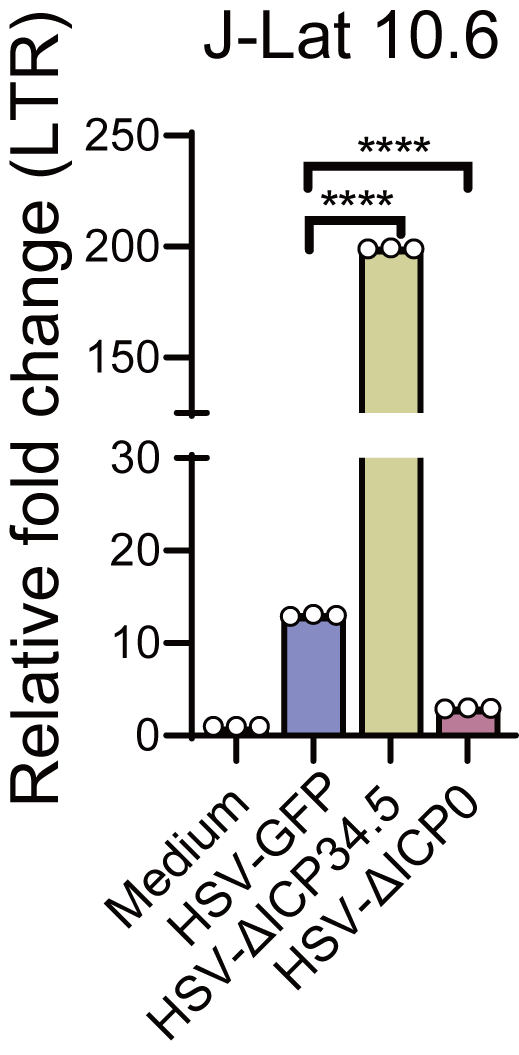

### Figure S5.tif

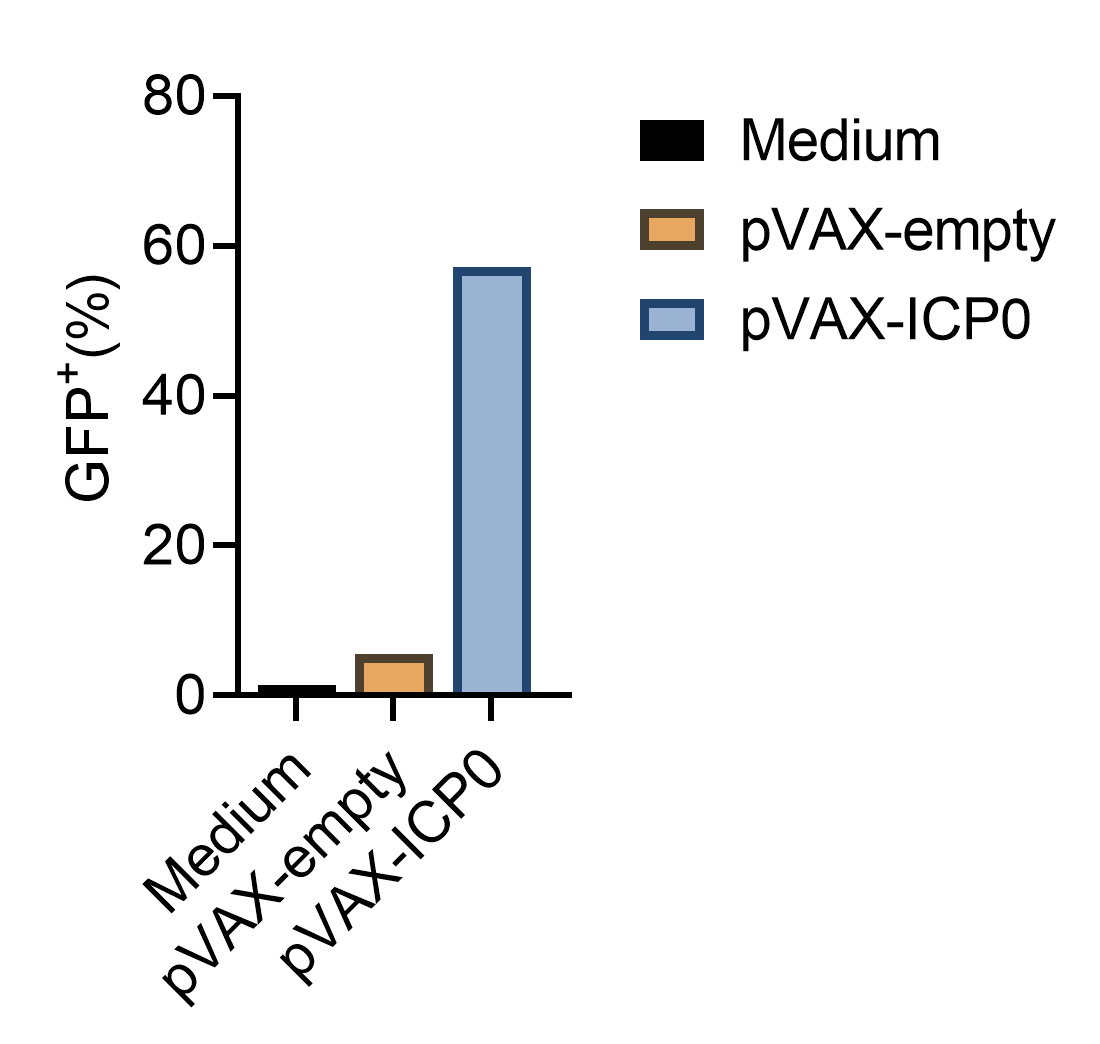

### Figure S6.tif

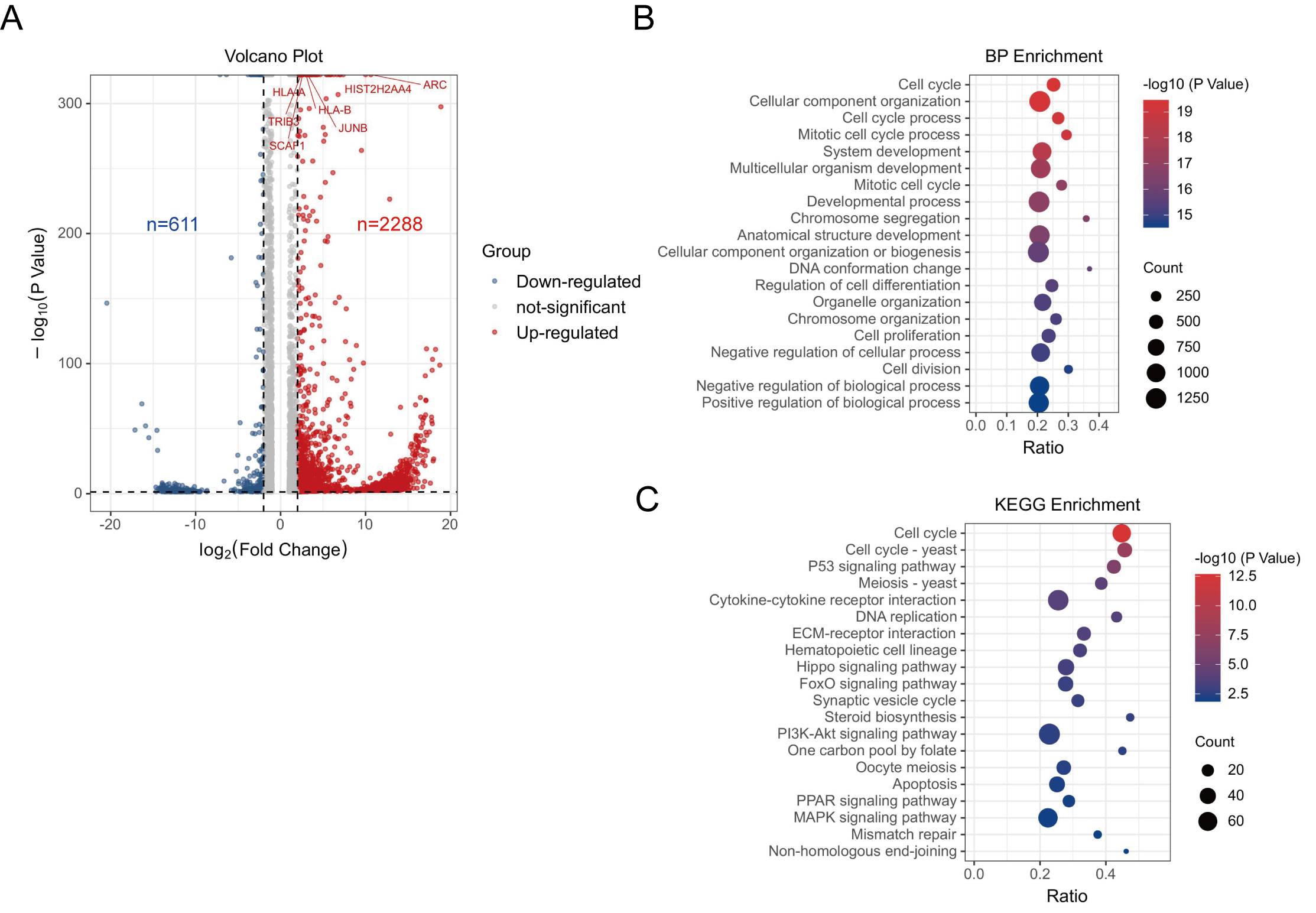

### Figure S7.tif

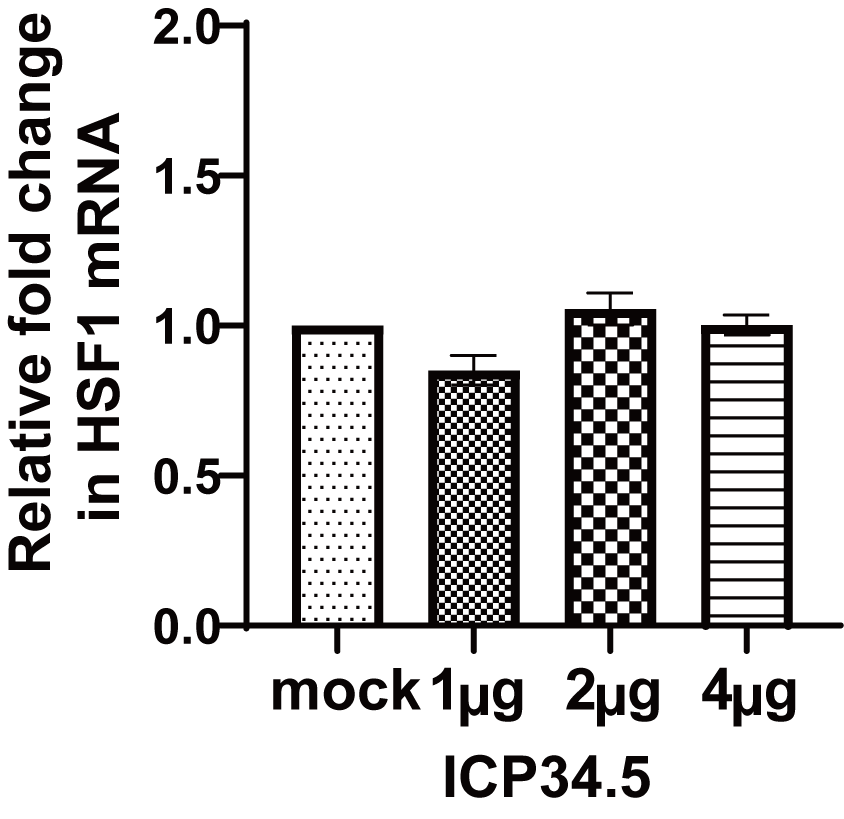

### Figure S8.tif

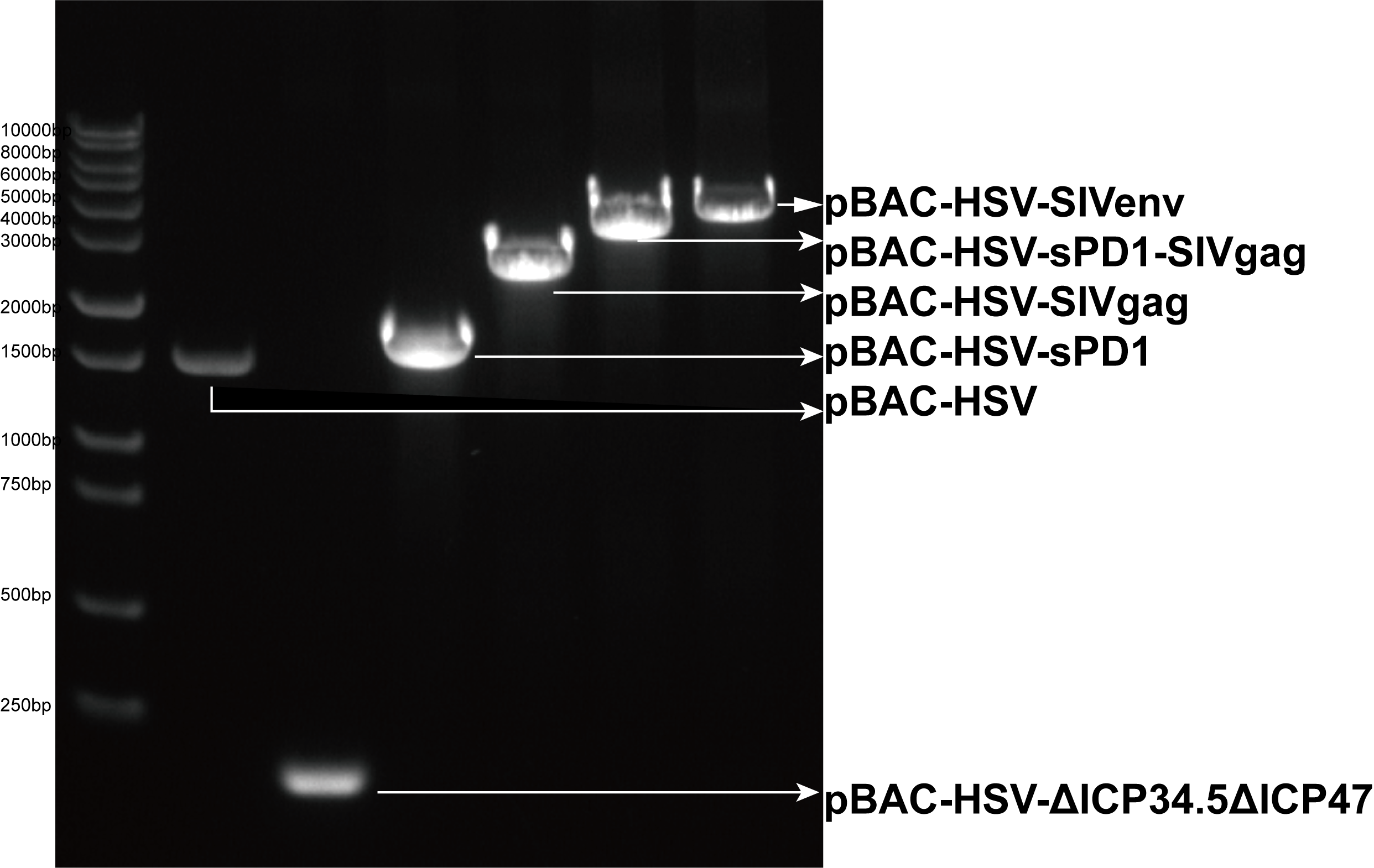

### Figure S9.tif

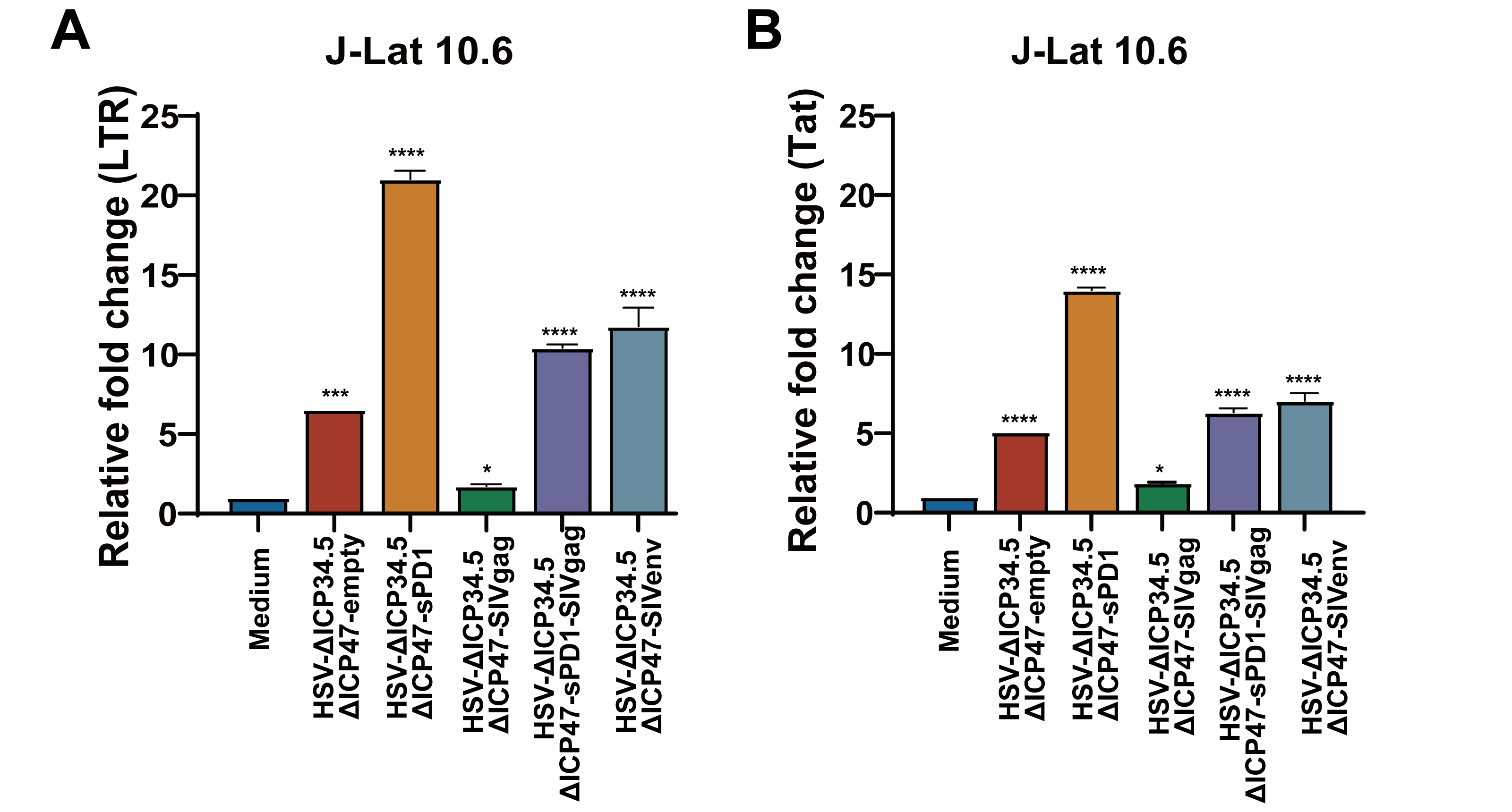

### Figure S10.tif

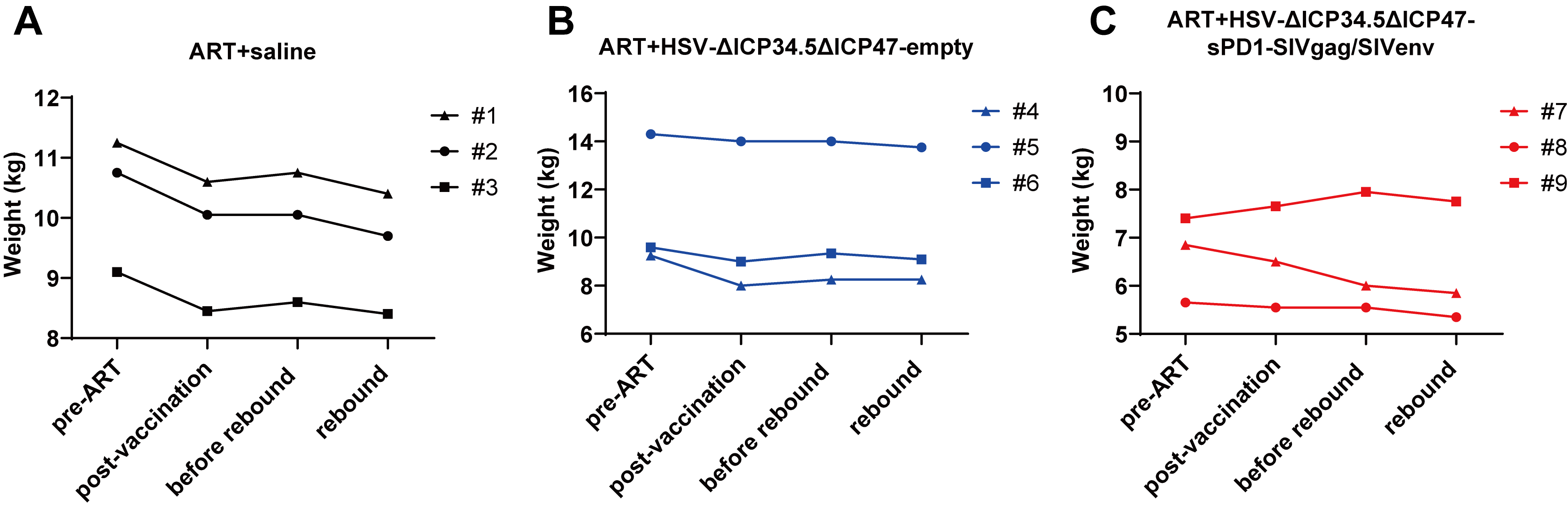

### Figure S11.tif

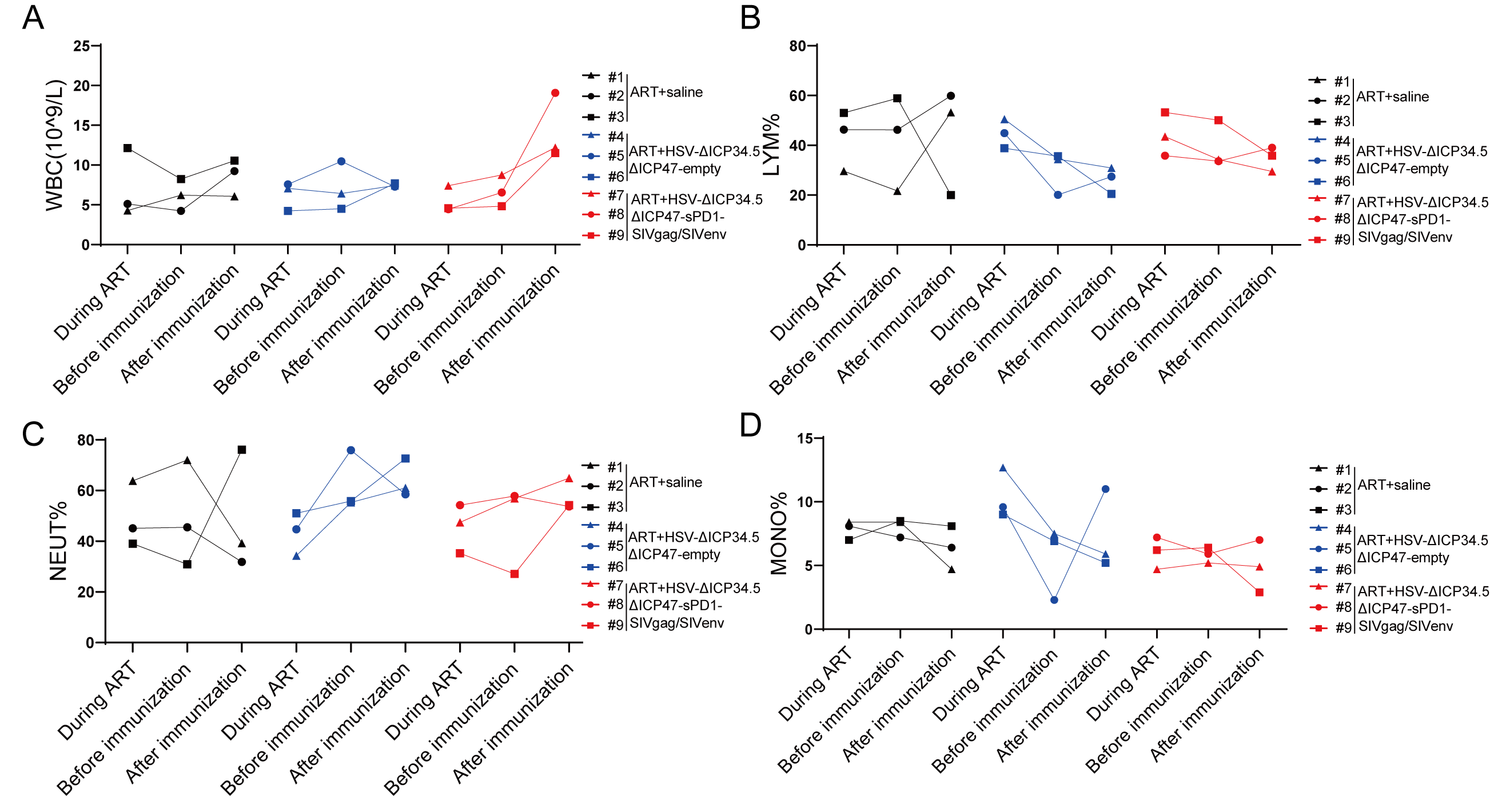
