## supplemental figures and tables for "Purging viral latency by a bifunctional HSV-vectored therapeutic vaccine in chronically SIV-infected macaques"

**Supplementary Information**

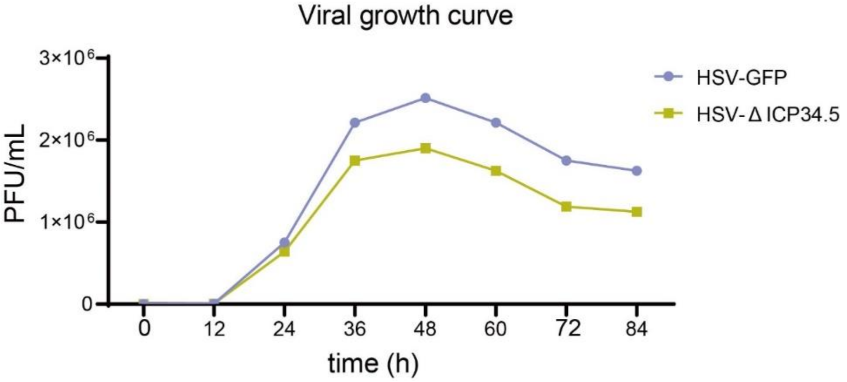


**Figure S1. Viral growth curve.** Vero cells were infected with HSV-GFP and HSV-ΔICP34.5 at an MOI of 0.01. Cells and supernatants were collected at various time points, and viral titers were determined by plaque assay.

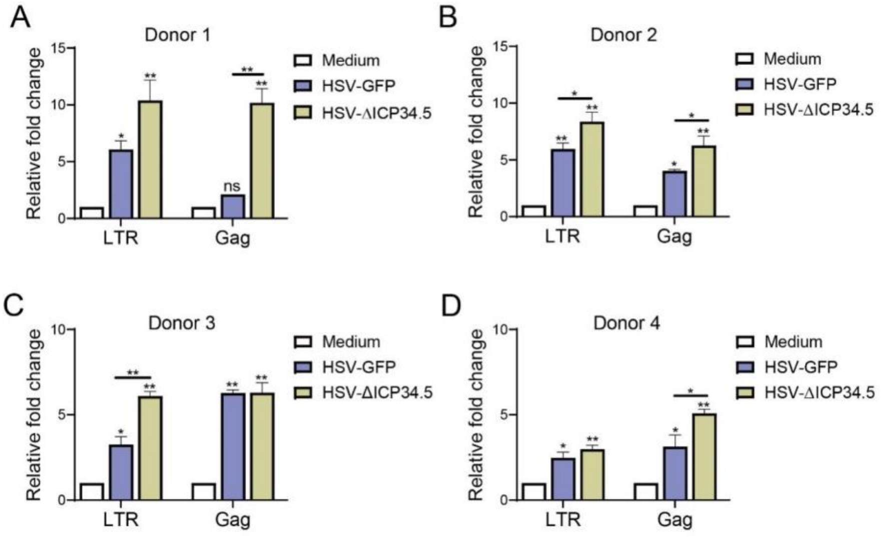


**Figure S2. The reactivation effect of HSV-1 on the latent HIV reservoir in primary CD4$^+$ T cells from PLWH.** The CD4$^+$ T cells from four donors of PLWH (A-D) were infected with HSV-GFP and HSV-ΔICP34.5, and then HIV-1 LTR, Gag were detected by quantitative PCR. Three independent experiments were repeated. **P*<0.05, ***P*<0.01. ns: no significance.

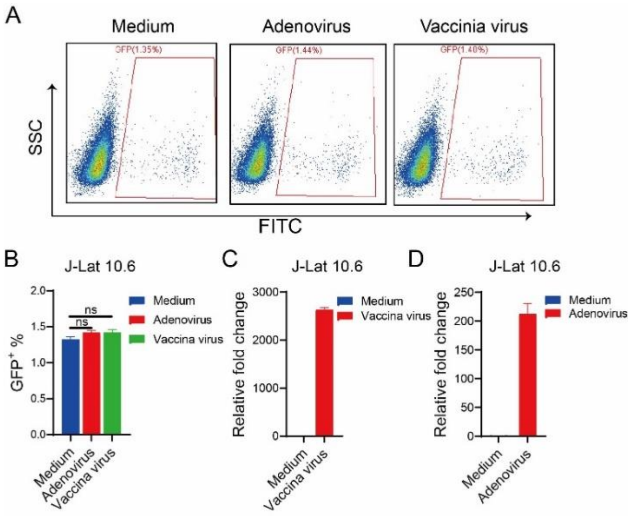


**Figure S3. Adenovirus and vaccinia virus cannot reactivate HIV latency in J-Lat 10.6 cells.** J-Lat 10.6 cells were infected with adenovirus and vaccinia virus for 30 h. The proportion of GFP+ cells, indicating activated latent cells, is shown in the pseudocolor plot (A) and the corresponding bar chart (B). The bar charts represent the mRNA levels of adenovirus L1 gene (C) and vaccinia virus tk gene (D).

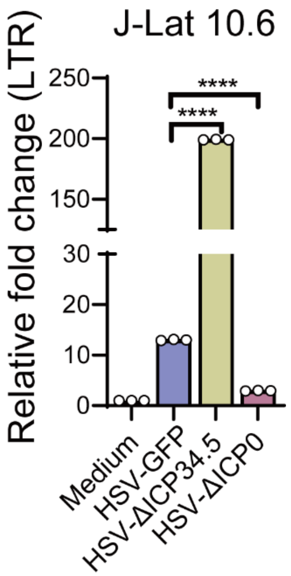


**Figure S4. Deletion of ICP0 diminished the reactivation effect of HIV latency by HSV-1.** The J-Lat 10.6 cells were infected with HSV-GFP, HSV-ΔICP34.5 and HSV-ΔICP0, and then HIV-1 LTR were detected by quantitative PCR. Three independent experiments were repeated. ****$P$<0.0001.

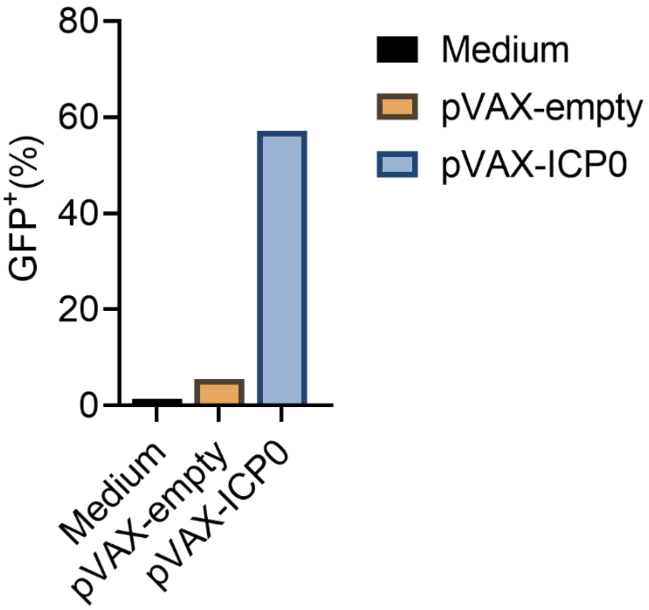


**Figure S5. ICP0 reactivates latent HIV in J-Lat 10.6 cells. J-Lat 10.6 cells (2 × $10^6$) were transfected with either pVAX-empty or pVAX-ICP0 for 24 h. The proportion of GFP⁺ cells, indicating activated latent cells, is shown in the bar chart.**

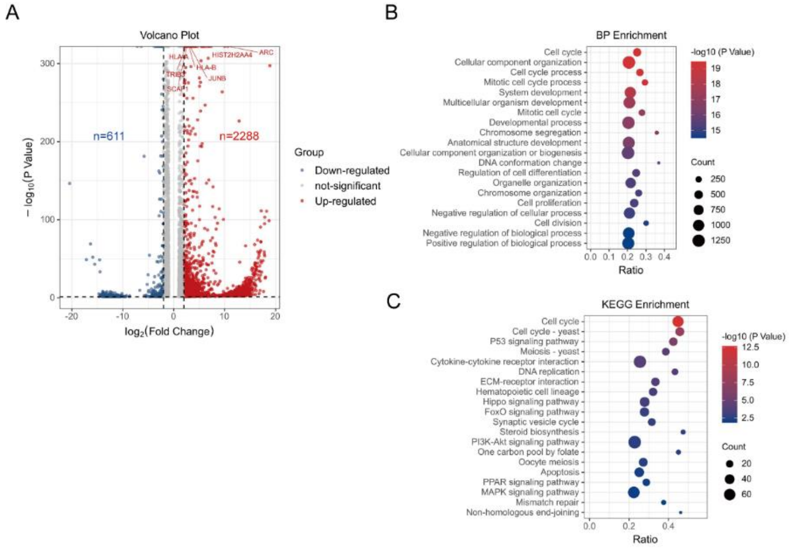


**Figure S6. RNA-seq analysis for HSV-ΔICP34.5-induced signaling pathways.** J-Lat 10.6 cells were either infected or not infected with HSV-GFP and HSV-ΔICP34.5, followed by extraction of total cellular RNA for sequencing and analysis. (A) Volcano plot showing differentially expressed genes between the two groups, with blue indicating downregulation and red indicating upregulation; (B) Bubble plot illustrating biological function enrichment analysis; (C) Bubble plot demonstrating enrichment of differential genes in signaling pathways. Circle size represents the number of genes enriched in that pathway, with colors ranging from blue to red indicating increasing -log10 (P-value) values.

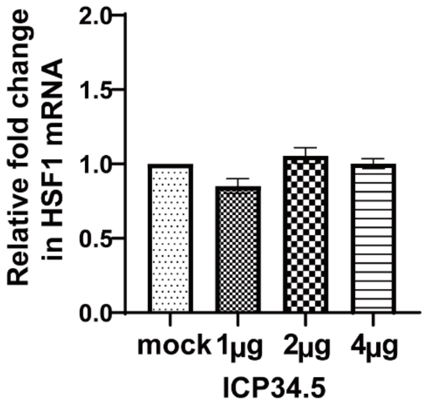


**Figure S7. The overexpression of ICP34.5 have no influence on the level of HSF1 expression.**

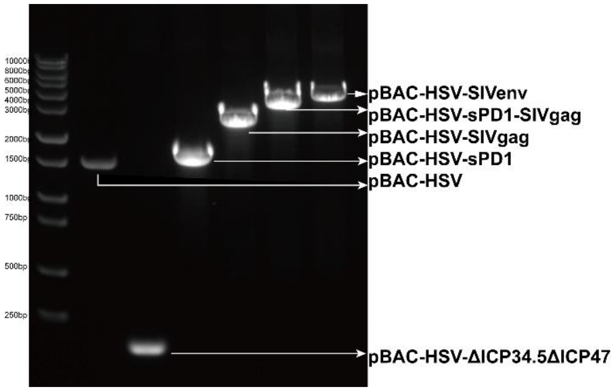


**Figure S8. The identification of recombinant pBAC-HSV plasmid.** Identification was performed by amplifying the pBAC plasmid using the tHSV-F and tHSV-R primers designed in the homologous arm of ICP34.5. lane1: DNA marker; lane2: pBAC-HSV (negative control); lane3: pBAC-HSV-∆ICP34.5∆ICP47; lane4: pBAC-HSV-sPD1; lane5: pBAC-HSV-SIVgag; Lane6: pBAC-HSV-sPD1-SIVgag; pBAC-HSV-SIVenv.

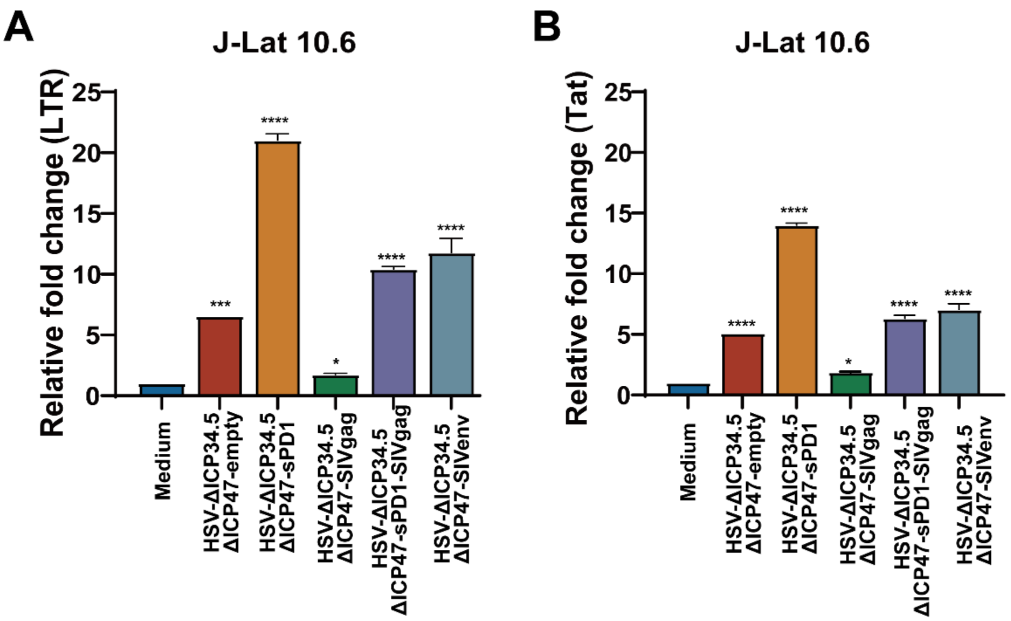


**Figure S9. The HSV-ΔICP34.5-based SIV vaccine effectively HIV latency.** J-Lat 10.6 cells ($1\times10^6$) were infected with recombinant HSV-empty, HSV-sPD1, HSV-ΔICP34.5ΔICP47-SIVgag, HSV-ΔICP34.5ΔICP47-sPD1-SIVgag, and HSV-ΔICP34.5ΔICP47-SIVenv, and then the relative fold changes of LTR (A) and Tat (B) were detected by quantitative PCR. *$P$<0.05, ***$P$<0.001, ****$P$<0.0001.

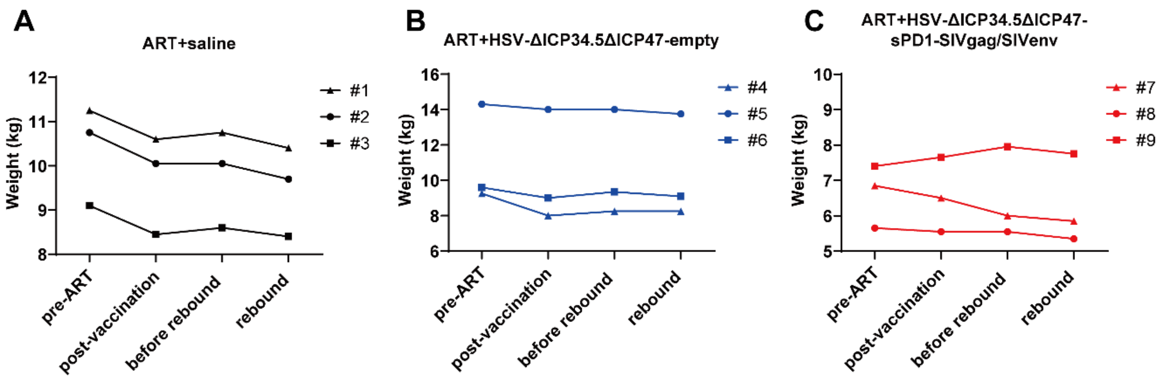


**Figure S10. The change of macaques' body weight during this study.**

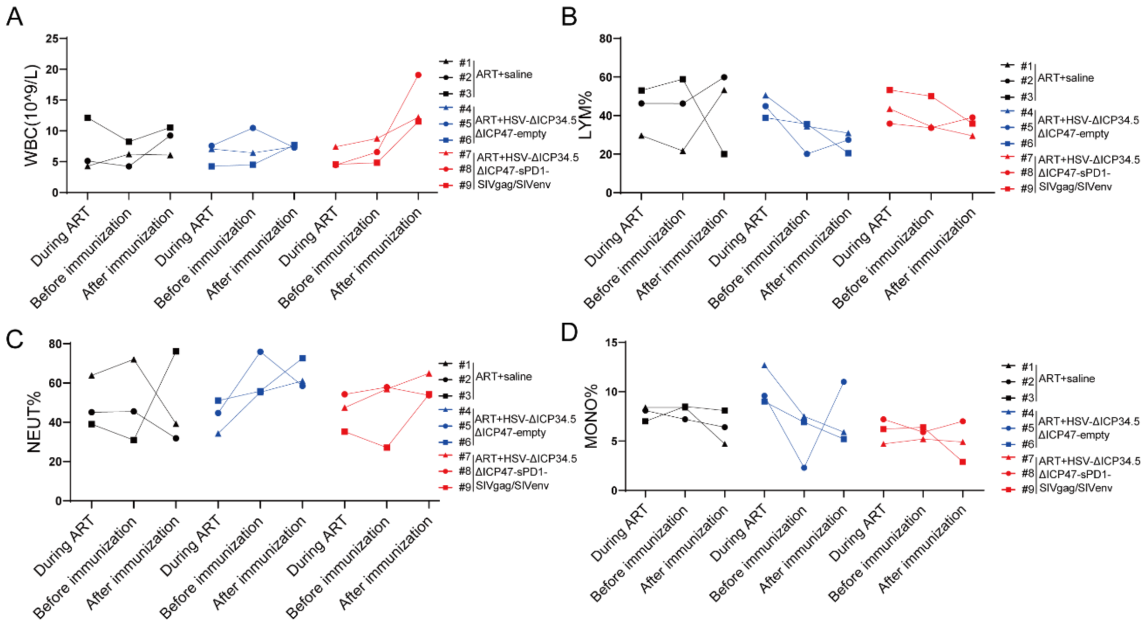


**Figure S11. Effect on the cell composition of peripheral blood in the macaques of different groups.** The levels of white blood cells (WBCs) (A), lymphocyte (LYM) (B), neutrophils (NEU) (C), and monocyte (MONO) (D) were monitored by the complete blood count tests.

**Table S1. Data for immunoprecipitation-mass spectrometry (IP-MS) analysis to identify potential molecules contributing to the HIV latency reactivation by HSV-ΔICP34.5-based constructs.**

| Accession | -10lgP | Avg. Mass |
|---|---|---|
| P0DMV8\|HS71A_HUMAN | 351 | 70052 |
| P11142\|HSP7C_HUMAN | 337.68 | 70898 |
| P10809\|CH60_HUMAN | 272.28 | 61055 |
| P34931\|HS71L_HUMAN | 261.48 | 70375 |
| P08238\|HS90B_HUMAN | 256.83 | 83264 |
| P34932\|HSP74_HUMAN | 202.19 | 94331 |
| P07900\|HS90A_HUMAN | 198.58 | 84660 |
| O95757\|HS74L_HUMAN | 146.39 | 94512 |
| Q92598\|HS105_HUMAN | 134.36 | 96865 |
| P04792\|HSPB1_HUMAN | 129.61 | 22783 |
| P62136\|PP1A_HUMAN | 186.52 | 37512 |

**Table S2. Baseline information for the experimental macaques in this study.**

| Number | Age (y) | Gender | Viral Load (Log10 copies/mL) | CD4 (count/mL) | Body Weight （kg） |
|---|---|---|---|---|---|
| #1 | 11 | M | 5.72 | 83 | 11.25 |
| #2 | 11 | M | 4.05 | 972 | 10.75 |
| #3 | 15 | M | 2.98 | 1819 | 9.10 |
| #4 | 19 | M | 4.35 | 835 | 9.25 |
| #5 | 11 | M | 5.57 | 238 | 14.3 |
| #6 | 11 | M | 3.24 | 599 | 9.60 |
| #7 | 11 | M | 5.99 | 1181 | 6.85 |
| #8 | 18 | F | 4.18 | 767 | 5.65 |
| #9 | 10 | F | 3.82 | 721 | 7.40 |

Note: y: years old; M: male; F: female.

**Table S3. Antibodies for the ICS assay in this study.**

| Classification | Antibodies |
|---|---|
| Mice experiment | Mouse CD3-FITC |
| | Mouse CD4-BB700 |
| | Mouse CD8-PECY7 |
| | Mouse IFN-$\gamma$-APC |
| | Mouse TNF-$\alpha$-PE |

| | |
|---|---|
| | Mouse IL-2-Violet605 |
| | Mouse CD44-Ko525 |
| | Mouse CD62L-BV786 |
| | Mouse CD107a-BV510 |
| | Mouse CD49b-PE |
| Monkey experiment | Hu/NHP CD4 FITC L200 |
| | Hu/NHP CD3 APC SP34-2 |
| | NHP CD45 PE D058-1283 |
| | CD8 Percp SK1 |
| | NHP CD3 BV650 SP34-2 |
| | Hu/NHP CD4 PE-CF594 L200 |
| | Hu/NHP CD8 APC-Cy7 RPA-T8 |
| | Hu/NHP IL-2 APC MQ1-17H12 |
| | Hu/NHP IFN-$\gamma$ PE 4S.B3 |
| | Hu/NHP TNF PE-Cy7 MAb11 |

**Table S4. The sequences of primes used in this study.**

| Primers | Sequences |
|---|---|
| Tat-F | ATGGAGCCAGTAGATCCTAG |
| Tat-R | TGCTTTGATAGAGAAGCTTG |
| LTR-F | GCCTCAATAAAGCTTGCCTTGA |
| LTR-R | TCCACACTGACTAAAAGGGTCTGA |
| Gag-F | GTCCAGAATGCGAACCCAGA |
| Gag-R | GTTACGTGCTGGCTCATTGC |
| Vpr-F | CCACAAAGGGAGCCATACAATG |
| Vpr-R | TTATGGCTTCCACTCCTGCC |
| Vif-F | CACACAAGTAGACCCTGACCT |
| Vif-R | CCCTACCTTGTTATGTCCTGCT |
| HSF1-F | CCACCTCCACCCCTGAAAAG |
| HSF1-R | GCACCAGCTGCTTCCCTGA |
| β-actin-F | TCACCAACTGGGACGAC |
| β-actin-R | TGTCACGCACGATTTCC |
| SYBR S mac-g238 S | AATACTGTCTGCGTCATCTGG |
| SYBR S mac-g382A | ATGGTGCTGTTGGTCTACTTG |
| tHSV-F | TGCTTGCCTGTCAAACTCTAC |
| tHSV-R | CCTCGGGTGTAACGTTAGAC |
| Ad-F | GAGTTGGCACCCCTATTCGA |
| Ad-R | GTTGCTGTGGTCGTTCTGGT |
| VVtk-F | ATAGATACGGAACGGGACT |

| | |
|---|---|
| VV-tk-R | TTGCCATACGCTCACAG |
